# Investigating Hybrid Deep Learning Architectures for Speech Envelope Reconstruction from EEG

**DOI:** 10.64898/2026.05.24.727471

**Authors:** Uday Sankar Gottipalli, Aditi Jha, Krishna Miyapuram

## Abstract

Reconstructing speech envelopes from electroencephalography (EEG) signals is a challenging but valuable task for brain-computer interfaces (BCIs), with applications in assistive communication for individuals with speech impairments. While deep learning has improved reconstruction accuracy, most existing approaches are restricted to single-layer architectures such as convolutional neural networks (CNNs). This limits their ability to capture the full complexity of spatio-temporal and structural EEG patterns. In this work, we systematically extend the VLAAI framework by evaluating 26 architectures that integrate CNNs, long short-term memory networks (LSTMs), and graph convolutional networks (GCNs) in both single-layer and hybrid configurations. Experiments on the 64-channel SparrKULee dataset demonstrate that CNNs remain the strongest standalone models, but hybrid designs—particularly CNN-LSTM and CNN-GCN-LSTM—achieve competitive or superior performance. These results highlight the importance of combining spatial, temporal, and graph-based processing, and provide practical guidelines for hybrid architecture design. Our study offers the first large-scale comparative analysis of hybrid models for EEG-based speech envelope reconstruction, advancing robust BCI systems for non-invasive speech decoding.

## I. Introduction

Brain-computer interfaces (BCIs) enable direct communication between the human brain and external systems, with applications in rehabilitation, assistive communication, and cognitive enhancement [1]. Reconstructing speech from electroencephalography (EEG) is one such task. The speech envelope represents temporal amplitude fluctuations, crucial for comprehension, carrying cues about words and phrases. EEG measures brain activity via scalp electrodes with high temporal resolution but are effected by artifacts, noise, and interference [15]. This makes envelope reconstruction task difficult, which leads to the requirement of advanced computational models.

Deep learning has shown strong potential for this task. The Very Large Augmented Auditory Inference (VLAAI) framework, based on Convolution Neural Networks (CNNs), achieved encouraging results. However, most of the prior models rely on a single network type. This limits performance, as CNNs capture only local spatial features, Long-Short Term Memory (LSTMs) model keeps tracks of temporal dependencies, and Graphical Convolutional Networks (GCNs) exploit structural relationships between EEG channels. Using only one of these restricts the ability to capture full spatio-temporal dynamics.

To address this, we extend VLAAI by systematically exploring CNN, LSTM, and GCN hybrids. We evaluated 26 architectures—ranging from single-layer baselines to multilayer hybrids—on the 64-channel SparrKULee dataset [12]. This comparison tests whether hybrids outperform single-layer models and identifies the most effective combinations for EEG-based speech envelope reconstruction.

## II. Background and Related Work

Speech envelope detection from EEG is an emerging topic in BCIs. Early methods relied on linear approaches such as canonical correlation analysis (CCA) [2], which maximized correlation between EEG features and stimulus envelopes. These methods confirmed feasibility but were limited in capturing nonlinear dynamics of brain responses, motivating the use of deep learning models.

### A. Single-Layer Architectures

Deep learning has been applied mainly through single-layer architectures. CNN have dominated because of their ability to extract spatial features from multi-channel EEG. The VLAAI framework [3], built on CNNs, achieved a median correlation of 0.15, improving over linear baselines (≈ 0.10). Tang et al. [4] also reported correlations up to 0.15 with an end-to-end CNN, while De Taillez et al. [5] reached 0.22 using dilated CNNs. Zhang et al. [6] explored CNN-MLP hybrids for subject-independent learning, showing CNNs’ adaptability. Recurrent networks, particularly LSTMs, have been explored less frequently for envelope reconstruction, but they naturally capture temporal dependencies. Broderick et al. [7] reported around 0.1 correlation using an LSTM, and Li et al. [8] showed their effectiveness in auditory attention detection (AAD) tasks, highlighting their potential to capture longrange temporal context.

GCNs represent a newer direction by modeling EEG electrodes as nodes in a graph, thus accounting for structural and spatial relationships beyond Euclidean distance. Though limited in envelope reconstruction, GCNs have shown strong performance in auditory attention and classification tasks [13], [14], suggesting that their ability to capture inter-channel dependencies could benefit envelope reconstruction as well.

### B. Hybrid Architectures

Single-layer networks each capture only one aspect of EEG signals: CNNs capture local spatial patterns, LSTMs handle temporal dynamics, and GCNs model uses structural relations. Hybrid architectures aim to combine these strengths.

Kawahara et al. [9] combined CNNs and LSTMs for dual-speaker AAD, reporting 85% accuracy, outperforming either network individually. Kim et al. [10] integrated a CNN encoder with a diffusion-based vocoder for reconstructing continuous speech from EEG, demonstrating improved overlap accuracy. Wang et al. [11] employed GCN-LSTM hybrids for unspoken bilingual speech tasks, outperforming LSTM-only baselines. Pahuja et al. [14] further demonstrated CNN-GCN ensembles improving AAD accuracy. Together, these studies confirm that hybrid designs capture complementary spatio-temporal and structural features more effectively than single-layer models.

### C. Research Gap

Despite these advances, most studies on EEG-based speech envelope reconstruction focus on CNN-only designs, with VLAAI [3] being a prime example. LSTMs and GCNs have been explored mainly in related auditory tasks rather than direct envelope reconstruction. Moreover, systematic evaluations of hybrid architectures remain limited. Particularly, combinations involving GCNs are underexplored. This motivates our work: a comprehensive comparison of 26 CNN, LSTM, and GCN-based architectures—both single and hybrid—for speech envelope reconstruction on the SparrKULee dataset.

## III. Dataset and Preprocessing

We used the SparrKULee dataset [12], which contains EEG recordings of speech-evoked responses from 85 young adults (18–30 years, normal hearing, native Dutch). Data from 78 subjects (publicly available) were analyzed. Recordings were made using a 73-channel EEG system while subjects listened to audiobook stimuli in 6–10 trials of 15 minutes each. The dataset is publicly available under a CC-BY-NC-4.0 license.

EEG and speech signals were temporally aligned with a 100–200 ms latency. Only 64 scalp channels were used. Data were segmented into 20-second windows at 64 Hz, bandpass filtered (0.5–40 Hz), and cleaned using ICA [16]. The speech envelope was extracted with a Hilbert transform, rectified, low-pass filtered (10–20 Hz), and downsampled to 64 Hz [17].

EEG windows were reshaped into 2D tensors (*C* × *T*), while speech envelopes were row vectors of length *T*.

## IV. Methodology

The proposed hybrid architecture for EEG-to-speech-envelope reconstruction integrates three core blocks: CNN, GCN, and LSTM. These blocks can be arranged in flexible sequences to form hybrid models. Input EEG data has shape (*N, T, C*), where *B* is batch size, *T* is time steps, and *C* is channels. Data is processed sequentially through a user-defined block sequence. Each block’s output is designed to match the input of the next block. A final linear layer maps the output to the target speech envelope shape (*B, T*). The model is trained end-to-end using Pearson correlation as both the loss and evaluation metric.

### A. CNN Block

The CNN block extracts local spatial and temporal features. It uses 1D convolutions along the time dimension to capture hierarchical patterns such as auditory-evoked potentials. This is effective for short-term dependencies in EEG related to speech. Each of the *p* layers consists of Conv1D, LayerNorm, Leaky ReLU, and optional ZeroPadding to maintain temporal length.

- **Input:** (*B, C*_in_, *T*)
- **Output:** (*B, C*_out_, *T*)

### B. LSTM Block

The LSTM block captures temporal dependencies in the EEG. It uses gated mechanisms to selectively store or forget information. This block has *q* LSTM layers, optionally bidirectional.

- **Input:** (*B, T, C*_in_)
- **Output:** (*B, T, C*_out_)

For a single LSTM cell at time *t*:

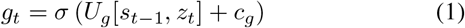

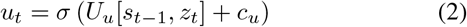

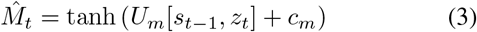

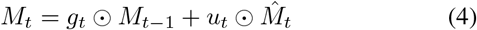

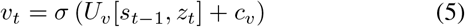

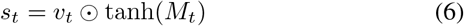

Here, 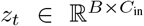represents the input vector at time step *t*, while *s*_*t−*1_ and *M*_*t−*1_ denote the previous hidden and memory states, respectively. The trainable weight matrices and bias terms are represented by *U*_*∗*_ and *c*_*∗*_, respectively. The sigmoid function *σ*(·) is used for gating operations, and ⊙ denotes element-wise multiplication. When *q* layers of LSTM cells are stacked, the combined output across all time steps is represented as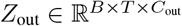.

### C. GCN Block

The GCN block models spatial relationships between EEG channels as a graph. Channels are nodes and edges represent inter-dependencies. It propagates information across nodes to capture global spatial patterns. The block has *r* GCN layers and uses a predefined adjacency matrix *G* ∈ R^*C×C*^. Each layer aggregates information from neighbors.

- **Input:** (*B, T, C*, 1)
- **Output:** (*B, T, C, f*)

A single graph convolutional layer can be formulated as:

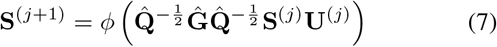

Here, 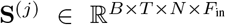 represents the input feature tensor at the *j*-th layer (initially, *F*_in_ = 1). The matrix **Ĝ** = **G** + **I** denotes the adjacency matrix augmented with self-loops, and is 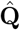 its corresponding degree matrix. The learnable parameter matrix is represented by **U**^(*j*)^, and *ϕ*(·) is a nonlinear activation function. Finally, a linear projection layer is applied to reshape the output back to the form (*B, T, N*).

### D. Hybrid Model Assembly and Training

The hybrid model (Fig. 1) is defined by a sequence of blocks, e.g., [CNN *p*, GCN *r*, LSTM *q*]. Block dimensions are aligned for compatibility; e.g., CNN output (*B, C*_out_, *T*) can be transposed to (*B, T, C*_out_) for LSTM input. The final linear layer predicts the envelope:

**Fig. 1.**
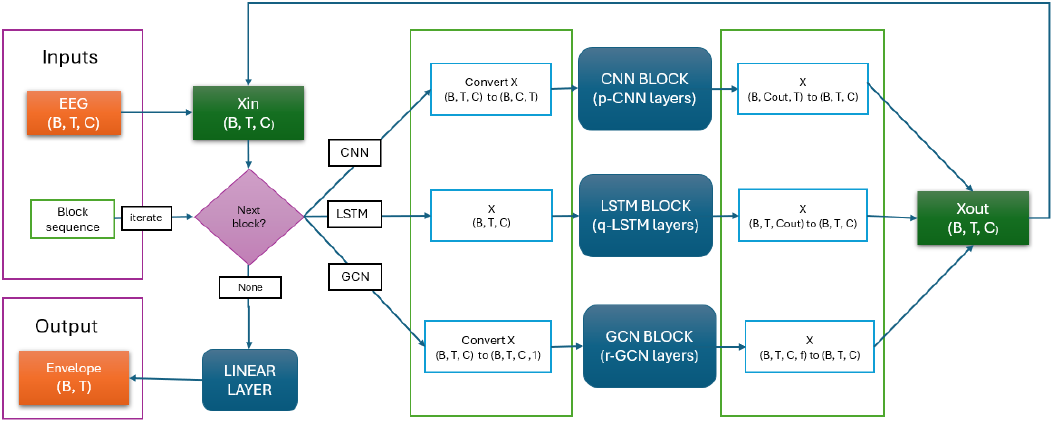
Architecture of the hybrid model for EEG-to-speech-envelope reconstruction. The model processes EEG input through a combination of CNN, GCN, and LSTM blocks using the block sequence, followed by a linear layer to produce the speech envelope. Here, *B* is Batch size, *T* is time steps, *C* is number of channels, where *Cin* and *Cout* specifies the input and output of a block, and *f* is features of each timestep of a channel

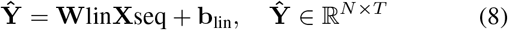

## V. Experimental Results

The Block sequence represents the structure of the architecture. The name of each model is a Block sequence that are given by combination of a number followed by a letter. Here, C implies CNN block, L implies LSTM block and G implies GCN block. And the number before the letter specifies the number of blocks of the specified type. Table I presents the test correlation performance for various hybrid model architectures used in EEG-to-speech-envelope reconstruction on the SparrKULee dataset [12]. Each architecture is defined by its Block sequence. The table show the mean and median correlation values of the models

The violin plots (fig. 2) shows the correlation values in the 26 hybrid models. The plot also shows the mean, median, and max in each model. The plot contains 5 subplots. A) Compares the single layer models of C and L. B) Compares the G-C and G-L hybrid models. C) Compares the C-L hybrid models. D) compares the G-C-L hybrid models where the first block is G, and E) compares the G-C-L hybrid models with the G block is in intermediate of the other blocks.

**Fig. 2.**
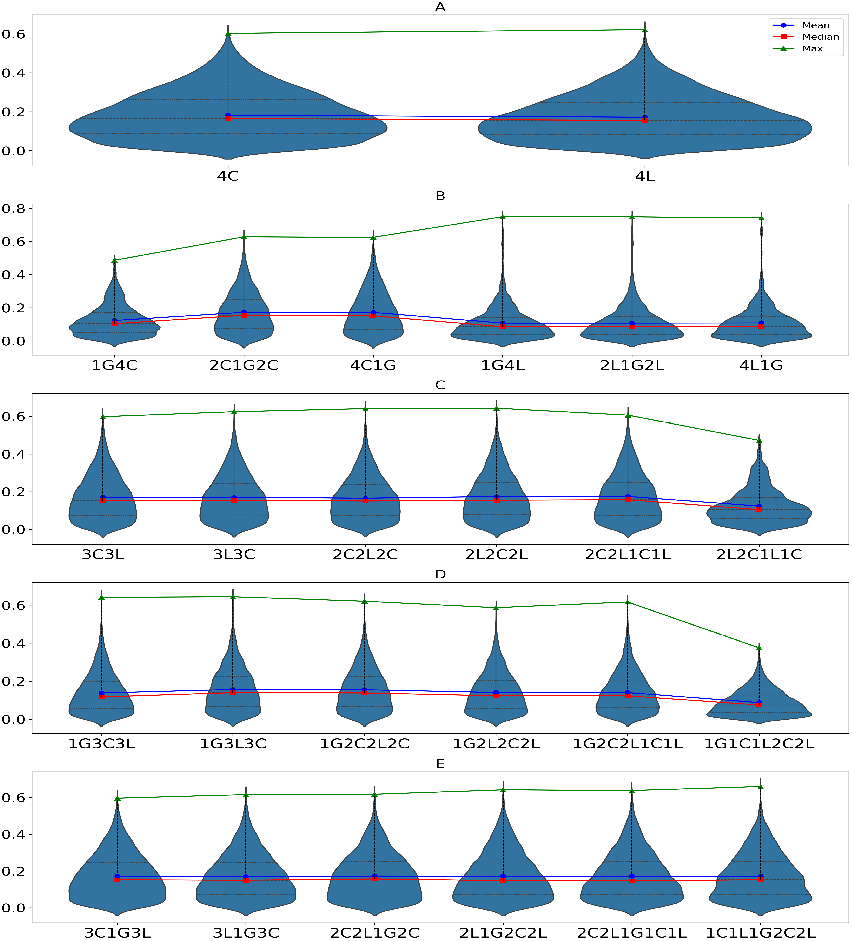
Violin plots along with the mean, median and max line plots for 26 hybrid models are shown in this figure. The name of the model is shown with letters C-CNN, L-LSTM and G-GCN. and the number previous to the letter specifies number of blocks. A) Compares the single layer models C and L. B) Compares the G-C and G-L hybrid models. C) Compares the C-hybrid models. D) compares the G-C-L hybrid models where the first block is G and E) Compares the G-C-L hybrid models with the G block is in middle of the other blocks.

## VI. Discussions

The experimental results in Table I and Fig. 2 highlight the performance of different architectures for EEG-to-speech-envelope reconstruction. The CNN-only model (4-CNN) achieved the highest performance, with a mean correlation of 0.1825 and a median of 0.1649. This reinforces previous findings from the VLAAI framework, while also showing that spatial filtering remains a critical component for robust reconstruction. Hybrid CNN-LSTM architectures, such as 2C2L1C1L (mean = 0.1745, median = 0.1571), performed competitively. This suggests that combining spatial and temporal processing can improve results.

GCN-LSTM hybrids underperformed, with the lowest performance in the 1G4L model (mean = 0.1073, median = 0.0857). LSTM alone can decode EEG patterns, but combined with GCN, performance drops. However, some GCN-LSTM models achieved high maximum correlations for specific samples, which are likely outliers. One contributing factor could be the reliance on predefined adjacency matrices, which may not fully reflect the dynamic channel relationships required for this task. Interestingly, the GCN-CNN-LSTM hybrids performed well when the GCN block was placed between the CNN and LSTM layers. This ordering appears to be important: CNNs first extract localized features, GCNs then capture channel relationships from these intermediate representations, and LSTMs finally model temporal dependencies. This finding provides valuable guidance for future hybrid model design, emphasizing that block sequence—not just block type—plays a decisive role in performance.

Overall, the results show that while CNNs remain the strongest individual architecture, carefully designed hybrids can approach or match their performance while offering additional flexibility. These findings help clarify when and how hybrid models should be deployed for EEG-based speech envelope reconstruction.

**Table 1.** Test correlation for different block architectures in EEG-to-speech-envelope reconstruction.

| Architecture (Block Sequence) | Mean Corr | Median Corr |
| --- | --- | --- |
| 4C | 0.1825 | 0.1649 |
| 4L | 0.1721 | 0.1553 |
| 1G4C | 0.1233 | 0.1040 |
| 2C1G2C | 0.1728 | 0.1547 |
| 4C1G | 0.1721 | 0.1509 |
| 1G4L | 0.1073 | 0.0857 |
| 2L1G2L | 0.1074 | 0.0848 |
| 4L1G | 0.1083 | 0.0853 |
| 3C3L | 0.1707 | 0.1522 |
| 3L3C | 0.1697 | 0.1541 |
| 2C2L2C | 0.1654 | 0.1504 |
| 2L2C2L | 0.1738 | 0.1548 |
| 2C2L1C1L | 0.1745 | 0.1571 |
| 1C1L2C2L | 0.1728 | 0.1527 |
| 1G3C3L | 0.1381 | 0.1072 |
| 1G3L3C | 0.1565 | 0.1397 |
| 1G2C2L2C | 0.1552 | 0.1383 |
| 1G2L2C2L | 0.1407 | 0.1230 |
| 1G2C2L1C1L | 0.1401 | 0.1228 |
| 1G1C1L2C2L | 0.1401 | 0.1227 |
| 3C1C3L | 0.1713 | 0.1518 |
| 3L1G3C | 0.1699 | 0.1497 |
| 2C2L1G2C | 0.1724 | 0.1566 |
| 2L1G2C2L | 0.1713 | 0.1473 |
| 2C2L1G1C1L | 0.1713 | 0.1457 |
| 1C1L1G2C2L | 0.1712 | 0.1521 |

## VII. Conclusion

This work makes three main contributions. First, it systematically evaluates 26 variants of CNN-, LSTM-, and GCN-based architectures, extending the VLAAI framework beyond CNN-only designs. Second, it provides the first large-scale comparison of hybrid combinations for EEG-to-speech-envelope reconstruction, offering insights into how spatial, temporal, and graph-based modeling interact. Third, it identifies effective design principles for hybrids, particularly the benefit of placing GCNs between CNNs and LSTMs to exploit complementary feature representations. The key takeaway is that CNNs remain the most reliable backbone for this task, but hybrid models, especially CNN-LSTM and CNN-GCN-LSTM can provide competitive or improved performance when appropriately structured. By clarifying the role of different architectural blocks and their combinations, this study advances our understanding of how to design deep learning systems for non-invasive BCIs.

Looking ahead, future research could focus on adaptive or learned adjacency matrices to enhance GCN performance, explore deeper or transformer-based hybrids, and validate findings across multiple EEG datasets. Finally, this work supports the creation of stronger BCI systems that helps in assistive communication by decoding speech from brain signals.

